# Fragile X Mental Retardation Protein modulates somatic D-type K^+^ channels and action potential threshold in the mouse prefrontal cortex

**DOI:** 10.1101/2020.07.30.228874

**Authors:** Brian E. Kalmbach, Darrin H. Brager

## Abstract

Axo-somatic K^+^ channels control action potential output in part by acting in concert with voltage-gated Na^+^ channels to set action potential threshold. Slowly inactivating, D-type K^+^ channels are enriched at the axo-somatic region of cortical pyramidal neurons of the prefrontal cortex where they regulate action potential firing. We previously demonstrated that D-type K^+^ channels are down regulated in extratelencephalic-projecting L5 neurons (ET) in the prefrontal cortex of the *fmr1* knockout mouse model of Fragile X syndrome (FX mice), resulting in a hyperpolarized action potential threshold. To test whether K^+^ channel alterations are regulated in a cell autonomous manner in FXS, we used a viral-mediated approach to restore expression of Fragile X Mental Retardation Protein (FMRP) in a small population of prefrontal neurons in male FX mice. Outside-out voltage clamp recordings revealed a higher D-type K^+^ conductance in FMRP-positive ET neurons compared to nearby FMRP-negative ET neurons. FMRP did not affect either rapidly inactivating A-type or non-inactivating K^+^ conductance. ET neuron patches recorded with FMRP_1-298_, a truncated form of FMRP which lacks mRNA binding domains, included in the pipette solution had larger D-type K^+^ conductance compared to heat-inactivated controls. Viral expression of FMRP in FX mice depolarized action potential threshold to near wild type levels in ET neurons. These results suggest that FMRP influences the excitability of ET neurons in the mPFC by regulating somatic D-type K^+^ channels in a cell autonomous, protein-protein dependent manner.

## INTRODUCTION

K^+^ channels are the most diverse voltage-gated ion channels present in neurons. Depending upon the localization and biophysical properties, K^+^ channels exert powerful control over action potential initiation, shape, and propagation. D-type channels are members of the *shaker* or K_V_1 family of voltage-gated K^+^ channels. D-type K^+^ channels inactivate with a time constant of hundreds of milliseconds, an order of magnitude slower than rapidly inactivating A-type K^+^ channels (Castellino et al., 1995; Jerng et al., 2004). This allows for a broader time window during which D-type channels can influence changes in membrane potential. D-type K^+^ channels are enriched in the axo-somatic region of neurons where they oppose depolarization and exert strong influence over the threshold for action potential generation (Kole et al., 2007; Higgs and Spain, 2011; Ordemann et al., 2019). Changes in K^+^ channel function contribute to neuronal hyperexcitability in neurological disorders such as epilepsy and episodic ataxia (Jan and Jan, 2012; Noebels et al., 2012; D’Adamo et al., 2020b). More recently, K^+^ channels have been implicated in neurodevelopmental disorders and learning and memory impairments (D’Adamo et al., 2020a).

One of these disorders, Fragile X syndrome (FXS), is the most common form of inherited mental impairment and the leading identified genetic cause of autism. Multiple classes of K^+^ channels are altered in mouse models of FXS. In hippocampal pyramidal neurons, loss of FMRP, the protein whose absence causes FXS, is associated with lower functional expression of A-type K^+^ channels (Gross et al., 2011; Routh et al., 2013) but see (Lee et al., 2011), and loss of Ca^2+^-activated BK and SK channel function (Deng et al., 2013; 2019). In the anterior ventral cochlear nucleus and medial nucleus of the trapezoid body, loss of FMRP reduces Na^+^-activated K^+^ channel (Slick/Slack) activity (Brown et al., 2010) and flattens the tonotopic map of K_V_3.1 channel expression (Strumbos et al., 2010). In L5 somatosensory neurons, loss of FMRP results in reduced dendritic I_BK_ (Zhang et al., 2014). Importantly, the loss of FMRP affects K^+^ channel function in a cell type specific manner. We demonstrated that extratelencephalic-projecting (ET) L5 neurons of the mPFC in FX mice have reduced somatic D-type K^+^ channel current (I_KD_), while neighboring intratelencephalic-projecting (IT) L5 neurons are unaffected (Kalmbach et al., 2015).

Here, we use sparse viral expression of FMRP in FX mice, in combination with electrophysiological recordings from visually identified FMRP-positive and FMRP-negative mPFC L5 pyramidal neurons, to test whether expression of FMRP rescues D-type K^+^ channel function. Viral expression of FMRP rescues D-type K^+^ channel function in L5 ET, but not IT, mPFC neurons. Intracellular perfusion through the recording electrode of a truncated FMRP, which lacks mRNA binding domains (Ramos et al., 2006), also rescues D-type K^+^ channel function, indicating that FMRP regulates D-type channel function via protein-protein interactions. Finally, we show that as a consequence of these alterations to K^+^ channels, action potential threshold is depolarized by FMRP expression.

## METHODS

### Stereotaxic injections

All procedures involving animals were performed with the approval of the University of Texas Animal Care and Use Committee. Male FX mice, 8 –12 weeks old, were anesthetized with isoflurane (1–4% mixed in oxygen), placed in a stereotaxic apparatus, and prepared for injections with craniotomies over the target injection regions. Injections were performed using a pulled glass pipette (10 –15 μm diameter tip) mounted on a Nanoject II small-volume injector (Drummond Scientific). Each individual injection was performed at a speed of 23 nL/s, separated by a 2–3 min interval. Recombinant adeno-associated viruses (rAAV)-expressing Cre-dependent FMRP, Cre-dependent tdTomato, and ER-Cre recombinase were injected into either into the PFC (AP: +1.5 mm; ML: ± 0.45 mm; DV: 2, 1.75, and 1.5 mm; 40 nl per location). Viral construction is described in detail in our recent paper (Brandalise et al., 2020). For all injections, the pipette was left in place for 3–5 min before removing it from the brain. Mice were given analgesics (carprofen; 5 mg/kg; TW Medical #PF-8507) post-surgery and monitored daily to ensure complete recovery. Mice received a single dose of tamoxifen (45 mg/kg; Sigma #H6278) one week post-surgery, to induce expression of FMRP and tdTomato. Tissue was prepared for electrophysiological recording 2 weeks post-tamoxifen.

### Slice preparation

Mice were anesthetized with a ketamine (100 mg/kg)/xylazine (10 mg/kg) cocktail and were perfused through the heart with ice-cold saline consisting of (in mM): 2.5 KCl, 1.25 NaH_2_PO_4_, 25 NaHCO_3_, 0.5 CaCl_2_, 7 MgCl_2_, 7 dextrose, 205 sucrose, 1.3 ascorbate and 3 sodium pyruvate (bubbled with 95% O_2_/5% CO_2_ to maintain pH at ∼7.4). A vibrating tissue slicer (Vibratome 3000, Vibratome Inc.) was used to make 300 μm thick coronal sections containing the mPFC. Slices were held for 30 minutes at 35°C in a chamber filled with artificial cerebral spinal fluid (aCSF) consisting of (in mM): 125 NaCl, 2.5 KCl, 1.25 NaH_2_PO_4_, 25 NaHCO_3_, 2 CaCl_2_, 2 MgCl_2_, 10 dextrose and 3 sodium pyruvate (bubbled with 95% O_2_/5% CO_2_) and then at room temperature until the time of recording.

### Electrophysiology

Slices were placed in a submerged, heated (32–34°C) recording chamber that was continually perfused (1−2 mL/minute) with bubbled aCSF containing (in mM): 125 NaCl, 3.0 KCl, 1.25 NaH_2_PO_4_, 25 NaHCO_3_, 2 CaCl_2_, 1 MgCl_2_, 10 dextrose, 3 sodium pyruvate, 0.025 D-APV, 0.02 DNQX, 0.005 CGP55845, and 0.002 gabazine. Slices were viewed either with a Zeiss Axioskop microscope and differential interference optics or a Zeiss AxioExaminer D microscope and Dodt contrast optics. Patch pipettes (4−8 MΩ) were pulled from borosilicate glass and wrapped with Parafilm to reduce capacitance.

### Outside-out voltage-clamp recordings

Outside-out recordings were made using an Axopatch 200B amplifier (Molecular Devices), sampled at 10 kHz, analog filtered at 2 kHz and digitized by an ITC-18 interface connected to a computer running Axograph X. The pipette solution contained (in mM): 120 K-gluconate, 16 KCl, 10 HEPES, 8 NaCl, 7 K_2_ phosphocreatine, 0.3 Na−GTP, 4 Mg−ATP (pH 7.3 with KOH). TTX (1 µM) was added to the extracellular saline. Three separate command protocols were used: 1 – depolarizing voltage commands (−70 to 70 mV in 20 mV steps) from a holding potential of −90 mV; 2 – depolarizing voltage commands following a 200-ms step to −22 to inactivate A-type K^+^ channels; and 3 - depolarizing voltage commands (−70 to 70 mV in 20 mV steps) from a holding potential of −22 mV to inactivate all transient K^+^ channels (Kalmbach et al., 2015). Activation data were fit to a single Boltzmann function using a least-squares program. Linear leakage and capacitive currents were digitally subtracted by scaling traces at smaller command voltages in which no voltage-dependent current was activated.

### Whole cell current clamp recordings

The pipette solution contained (in mM): 120 K-gluconate, 16 KCl, 10 HEPES, 8 NaCl, 7 K_2_ phosphocreatine, 0.3 Na−GTP, 4 Mg−ATP (pH 7.3 with KOH). Neurobiotin (Vector Laboratories; 0.1-0.2%) was also included for histological processing and post-hoc cell location determination. In some cases, Alexa 594 (16 μM; Invitrogen) was also included in the internal recording solution to determine the dendritic recording location relative to the soma. Data were acquired using a Dagan BVC−700 amplifier (Dagan Inc.) and custom data acquisition software written using Igor Pro (Wavemetrics) or AxoGraph X (AxoGraph Scientific) data acquisition software. Data were acquired at 10−50 kHz, filtered at 2−10 kHz and digitized by an ITC-18 (InstruTech) interface. Pipette capacitance was compensated for and the bridge balanced during each recording. Series resistance (10−25 MΩ) was monitored and compensated throughout each experiment. Voltages are not corrected for the liquid-junction potential (estimated as ∼8 mV).

Data were analyzed using either custom analysis software written in Igor Pro or using AxoGraph X. Single action potentials (APs) were elicited using just-threshold current injections of various durations. AP threshold was defined as the voltage where the first derivative first exceeded 20 mV/ms.

### Data Analysis

Repeated measures analysis of variance (RM−ANOVA), between-subjects factors ANOVA, mixed factors ANOVA and post−hoc t−tests were used to test for statistical differences between experimental conditions. Sidak’s correction was used to correct for multiple comparisons. Error bars represent standard error of measurement (SEM). Statistical analyses were performed using Prism (Graphpad). Data are presented in the text as mean ± SEM.

## RESULTS

### Sparse expression of FMRP in the mPFC of FX mice

FMRP and the fluorescent reporter td-Tomato were expressed in a sparse population of L5 mPFC neurons in FX mice using an inducible viral expression system (Brandalise et al., 2020) (Figure 1). We recently demonstrated that this approach results in coexpression of tdTomato and FMRP, thereby allowing the comparison of FMRP+ (tdTomato+) and FMRP- (tdTomato-) neurons in the same slice (Brandalise et al., 2020). Furthermore, sparse expression of FMRP is effective in rescuing dendritic h-channel (I_h_) function in L5 ET neurons (Brandalise et al., 2020). Whether K^+^ channel function can be similarly rescued is not known.

**Figure 1.**
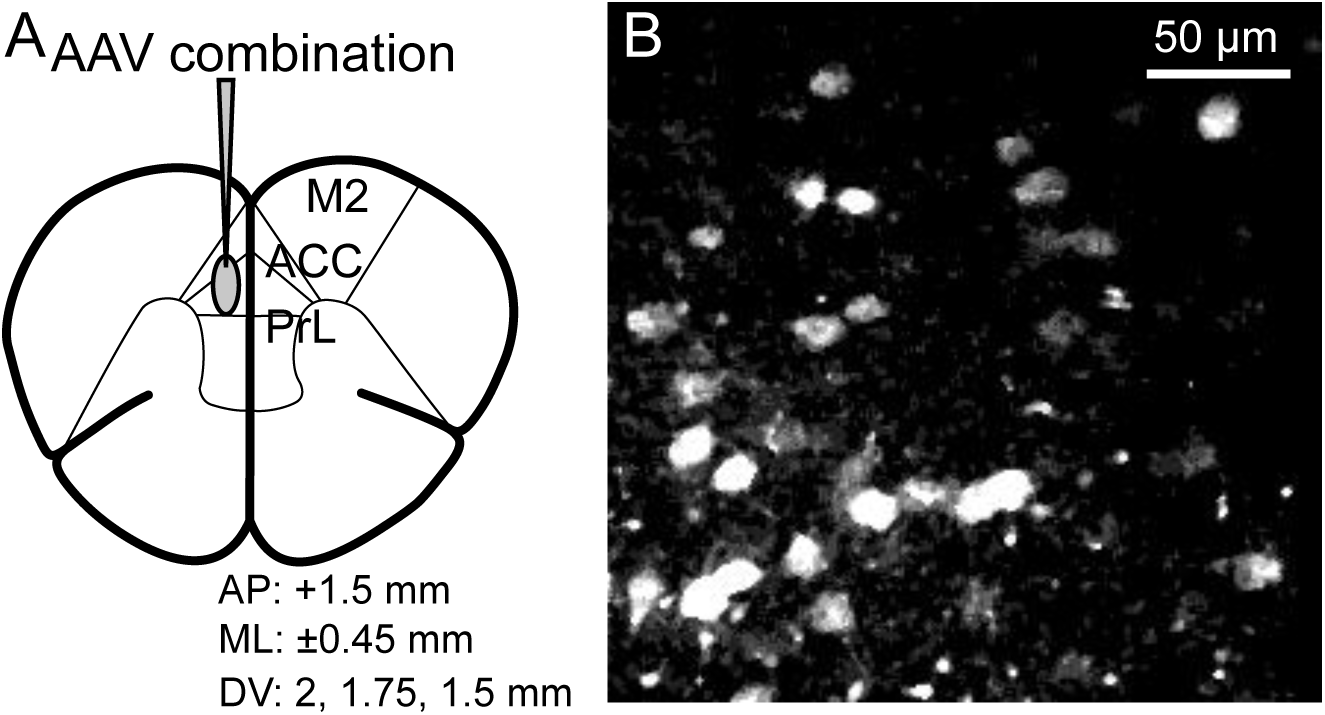

### FMRP expression decreases I_KD_ in ET neurons

To test whether FMRP regulates K^+^ currents in a cell autonomous manner, we used outside-out patch clamp recordings to measure K^+^ currents from FMRP+ and FMRP-L5 mPFC neurons. The outside-out configuration allowed us to identify the neuron as ET vs. IT based on intrinsic resonance prior to switching to voltage-clamp (Dembrow et al., 2010; Kalmbach et al., 2015). As previously demonstrated (Kalmbach et al., 2015), the total somatic I_K_ in ET and IT neurons could be separated into three components: a rapidly-inactivating A-type current (I_KA_), a slowly-inactivating current (I_KD_), and a sustained current (I_K-sust_). There was no significant difference in I_KA_ (unpaired t-test; t=0.3724, df=13, p=0.7156) and I_K-SUST_ (unpaired t-test; t=0.8146, df=13, p=0.4300) between FMRP+ and FMRP-ET neurons (Fig. 2A, C). We found however, that I_KD_ was significantly greater in FMRP+ compared to FMRP-ET neurons (Fig. 2B-C; unpaired t-test; t=4.498, df=13, p=0.0006). When normalized for patch area, D-type K^+^ conductance density was significantly higher in FMRP+ compared to FMRP-ET neurons (Fig. 2D; unpaired t-test; t=4.788, df=13, p=0.0001). A-type and sustained K^+^ conductance density was not different between FMRP+ and FMRP-patches. Expression of FMRP did not affect the voltage-dependence of I_KA_ or I_KD_ activation (Fig. 2E-F; Table 1). We previously showed that there was no difference in I_KD_ in L5 IT neurons between wild type and FX mice (Kalmbach et al., 2015). In agreement, we found no significant differences in I_KA_ (unpaired t-test; t=0.9865, df=6, p=0.3620) or I_KD_ (unpaired t-test; t=0.1630, df=6, p=0.8784) between FMRP+ and FMRP-IT neurons (Fig. 3A-C). When normalized for patch area, there was no significant difference in A-type (unpaired t-test; t=0.5207, df=6, p=0.6212) and D-type (unpaired t-test; t=0.1682, df=6, p=0.8720) K^+^ conductance density (Fig. 3D). These results suggest that expression of FMRP in the adult mPFC of FX mice increases D-type K^+^ channel conductance in L5 ET but not IT neurons.

**Figure 2.**
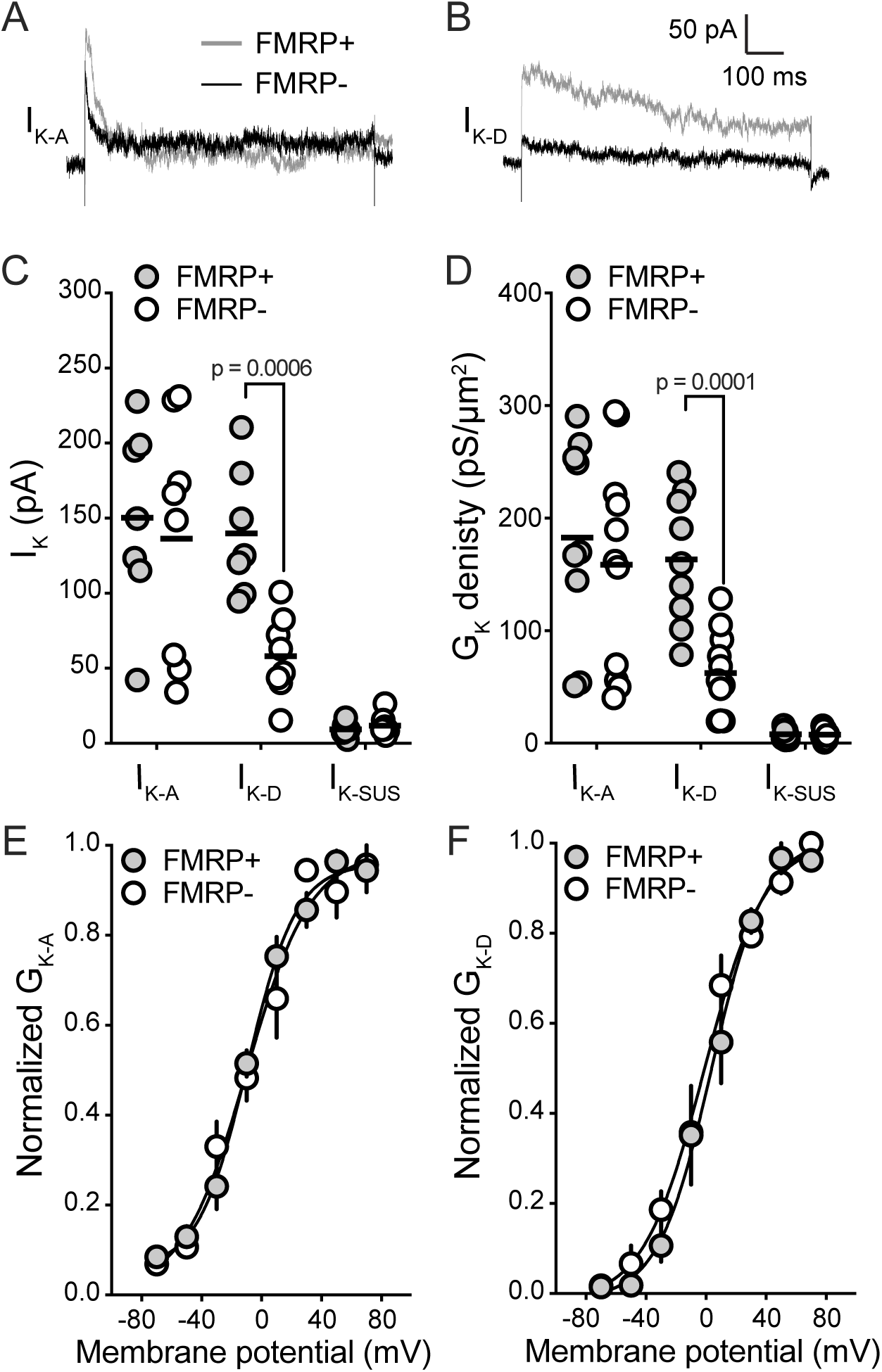

**Figure 3.**
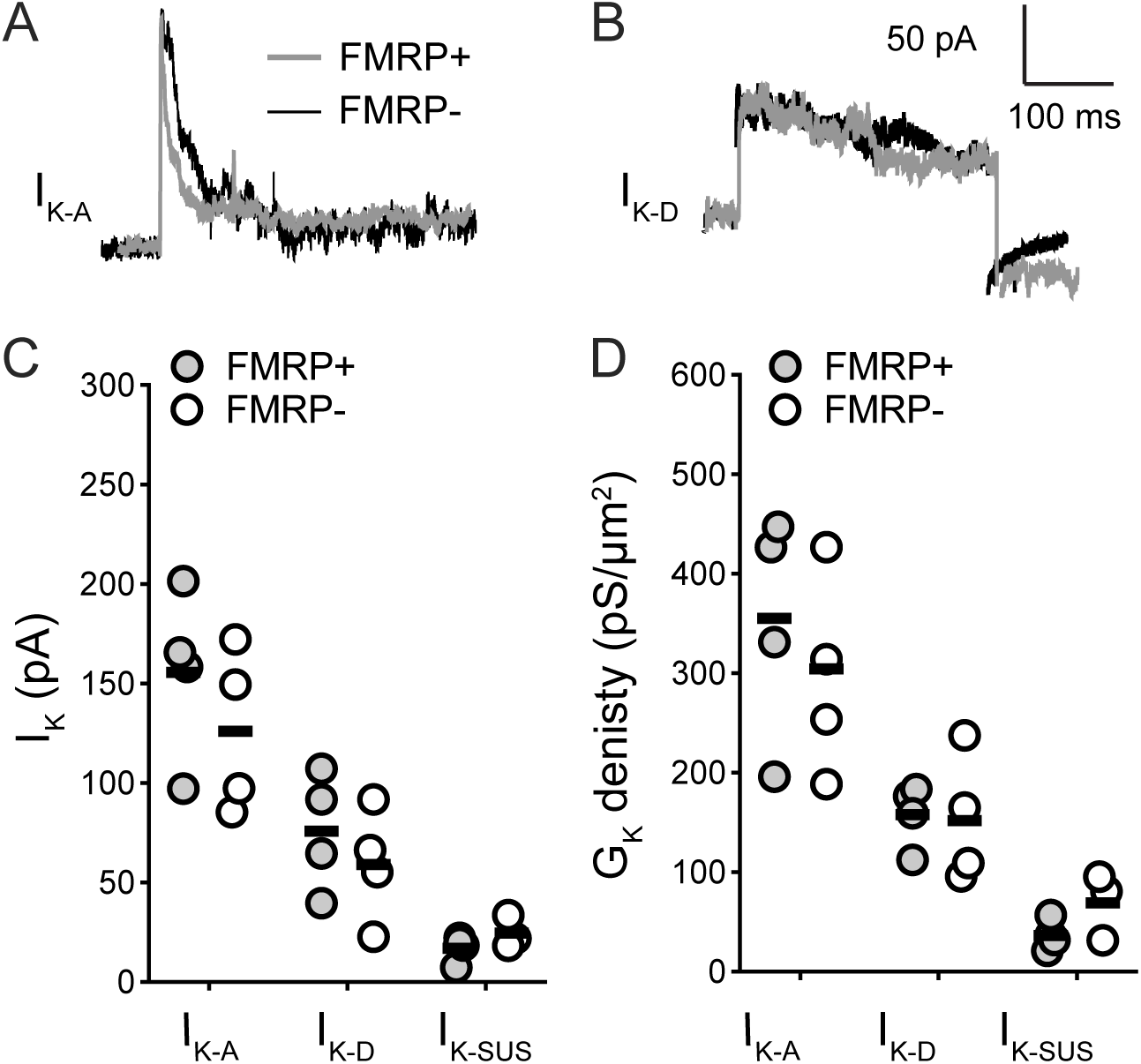

### FMRP regulates in ET neuron D-type K^+^ channels via protein-protein interactions

FMRP is traditionally considered to be a translational regulator which can bind to target mRNAs and repress or promote translation (Darnell et al., 2011). FMRP also has protein-protein binding domains in the N-terminal region through which FMRP can bind to and regulate protein function (Ramos et al., 2006). A truncated form of FMRP containing the N-terminal domain, FMRP_1-298_, lacks the mRNA binding domains but retains the ability to bind via protein-protein interactions (Ramos et al., 2006). FMRP_1-298_ interacts with and modulates Na^+^-activated and Ca^2+^-activated K^+^ channels (Brown et al., 2010; Deng et al., 2013; 2019). We recently used FMRP_1-298_ to demonstrated that FMRP regulates dendritic h-channels via protein-protein interactions in as little as 3 minutes (Brandalise et al., 2020).

To determine whether FMRP similarly regulates D-type K^+^ channel function, we made outside-out voltage-clamp measurements with either FMRP_1-298_ or heat-inactivated (HI) FMRP_1-298_ the pipette recording solution. We first measured cell resonance in current clamp, before pulling patches for voltage clamp recording, ensuring sufficient time for perfusion of FMRP_1-298_. I_KA_ and I_K-SUST_ were not different between patches recorded with FMRP_1-298_ and HI FMRP_1-298_ (Fig. 4A, C). Similar to our experiments using viral expression of full length FMRP, patches with FMRP_1-298_ had significantly higher I_KD_ compared to patches recorded HI FMRP_1-298_ (Fig. 4B-C; unpaired t-test; t=5.911, df=12, p<0.0001). D-type K^+^ conductance density was similarly significantly higher with FMRP_1-298_ compared to HI FMRP_1-298_ (Fig. 4D; unpaired t-test; t=3.737, df=12, p=0.0025). A-type and sustained K^+^ current and conductance density were not affected by FMRP_1-298_. Consistent with our viral results, FMRP_1-298_ did not significantly affect the voltage-dependence of activation for I_KA_ or I_KD_ in ET neurons (Fig. 4E-F; Table 2).

**Figure 4.**
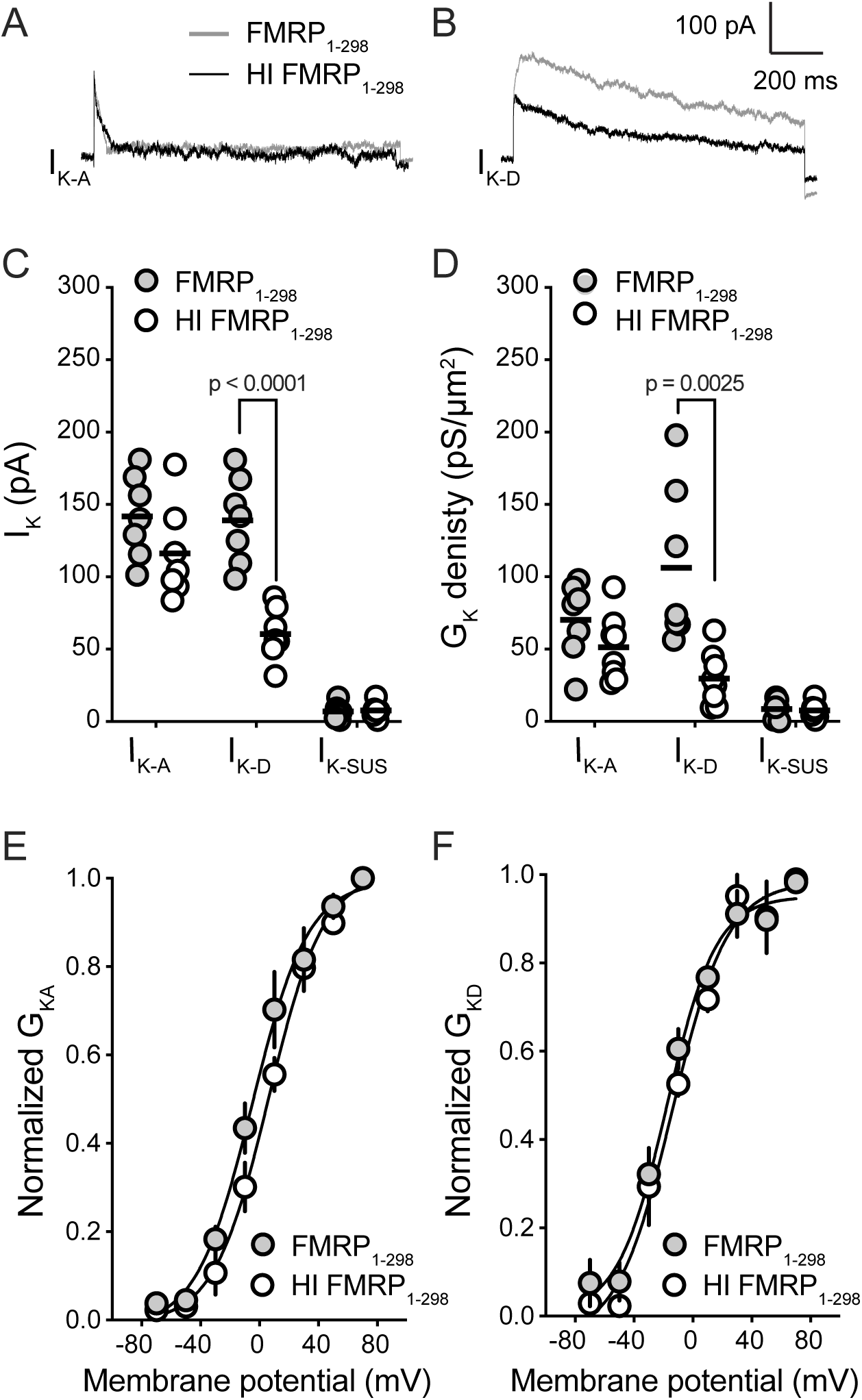

We recently showed that FMRP_1-298_ increased dendritic I_h_ in ET neurons of FX mice at concentrations as low as 10 nM (Brandalise et al., 2020). To test if lower concentrations of FMRP_1-298_ were also effective at increasing I_KD_, we repeated the above outside-out patch experiments with 10 nM FMRP_1-298_ in the pipette recording solution. We found that I_KD_ was significantly higher in outside-out patches from ET neurons recorded with 10 nM FMRP_1-298_ compared to HI FMRP_1-298_ (FMRP_1-298_: 162 ± 13.3 pA; HI FMRP_1-298_: 64 ± 12.8 pA; unpaired t-test; t=5.340, df = 6; p=0.0018). Given that the application of FMRP in these experiments is acute, these results suggest that FMRP regulates D-type K^+^ channel function via cell autonomous, protein-protein dependent mechanisms.

### Viral expression of FMRP depolarizes AP threshold in L5 ET neurons

Finally, we asked if expression of FMRP rescues action potential threshold in L5 PFC neurons in FX mice. We made whole-cell current clamp recordings from FMRP+ and FMRP-L5 neurons three weeks after stereotaxic delivery of viruses in FX mice. ET and IT neurons were identified based on their resonant profiles (Dembrow et al., 2010; Kalmbach et al., 2015). The biophysical properties of D-type K^+^ channels allow them to act as high pass filters by depolarizing the voltage threshold for action potential generation in response to slow, but not fast, voltage trajectories. It was previously shown that current injections of variable duration can be used to separate the contribution of I_KD_ to action potential threshold (Higgs and Spain, 2011; Kalmbach et al., 2015; Ordemann et al., 2019). Given that FMRP increases I_KD_ in L5 ET, we expect that expression of FMRP will depolarize ET neuron action potential threshold for longer current injections. Indeed, we found that action potential threshold was significantly depolarized in FMRP+ ET neurons compared FMRP-ET neurons (Fig. 5 A-B; Mixed factor ANOVA, interaction between FMRP and duration – F(6, 192)=2.220, p=0.0432). Consistent with higher I_KD_, post hoc tests revealed that action potential threshold was significantly depolarized for current injections longer than 12 ms (Kalmbach et al., 2015). Action potential threshold in FMRP+ ET neurons was not significantly different from action potential threshold in wild type ET neurons. As expected, expression of FMRP had no effect on action potential threshold in L5 IT neurons (Fig. 5 C-D; Mixed factor ANOVA, main effect of FMRP – F(1, 12)=0.09807, p=0.7595). The effect on action potential threshold in L5 ET neurons was specific to FMRP as expression of tdTomato alone did not affect action potential threshold (Fig. 5 E-F; Mixed factor ANOVA, main effect of tdTomato – F(1, 17)=0.02550, p=0.8750). These results suggest that expression of FMRP in the mPFC of adult FX mice restores action potential threshold to wild type levels in L5 ET neurons.

**Figure 5.**
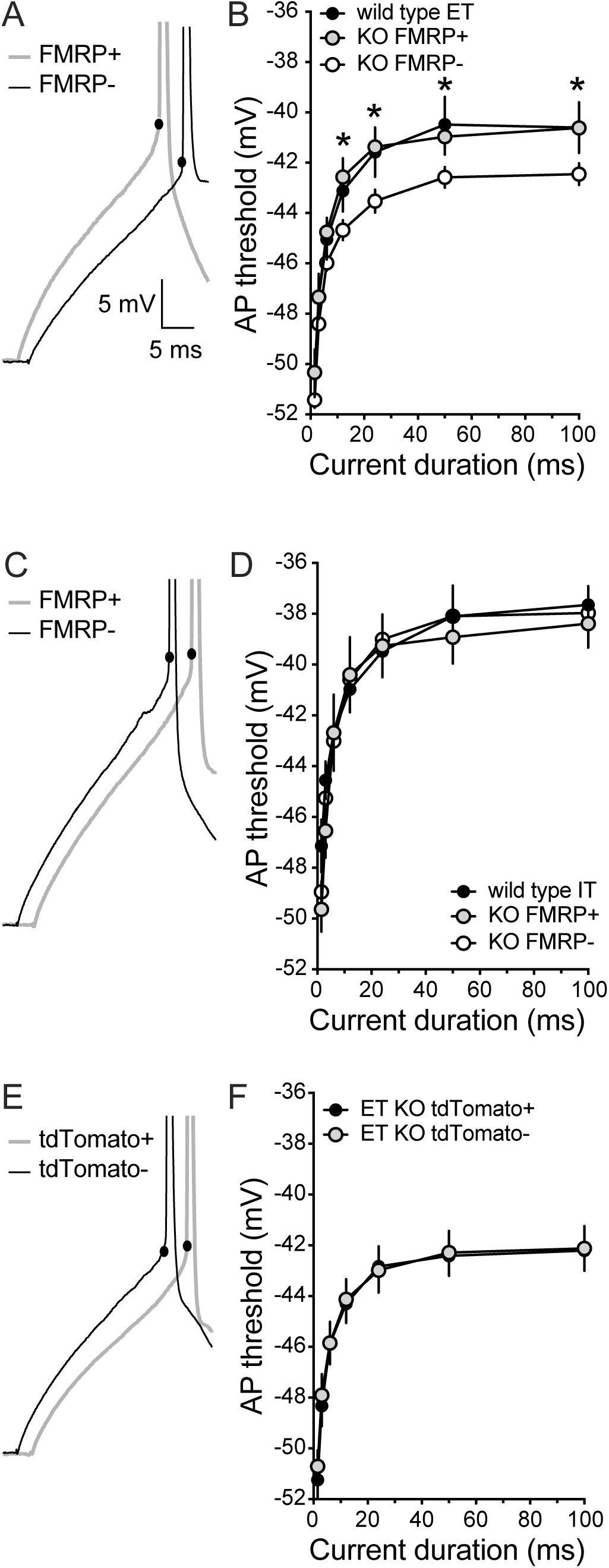

## DISCUSSION

K^+^ channels are critically important for regulating neuronal function. Dendritic K^+^ channels control the amplitude and duration of local synaptic inputs and back propagating action potentials (reviewed in (Johnston et al., 2003). Somatic K^+^ channels regulate the threshold and repetitive firing pattern of action potentials (Wang et al., 1998; Shao et al., 1999; Gu et al., 2007; Guan et al., 2007; Higgs and Spain, 2011; Guan et al., 2018). Many of these K^+^ channels are regulated by FMRP and thus altered in Fragile X syndrome including: rapidly inactivating, A-type channels (Gross et al., 2011; Lee et al., 2011; Routh et al., 2013; Kalmbach et al., 2015; Routh et al., 2017), Ca^2+^ activated channels (Deng et al., 2013; Zhang et al., 2014; Deng et al., 2019), Na^+^-activated channels (Brown et al., 2010; Zhang et al., 2012), and delayed rectifier channels (Strumbos et al., 2010; El-Hassar et al., 2019). FMRP regulates the function and expression of these K^+^ channels through translational and protein-protein interaction mechanisms. We previously showed that slowly inactivating D-type K^+^ channels are downregulated in L5 ET neurons of the mPFC (Kalmbach et al., 2015). In this study, we demonstrate that restoration of FMRP expression via viral delivery to the mPFC upregulates D-type K^+^ channel function and rescues action potential threshold in ET, but not IT, neurons of the mPFC. Furthermore, we demonstrate the FMRP regulates D-type K^+^ channels via protein-protein interactions as intracellular perfusion of FMRP_1-298_ was sufficient to rescue I_KD_.

### Possible mechanisms

We found that FMRP increased D-type K^+^ conductance but did not affect the voltage dependence of D-type K^+^ channel activation, in ET prefrontal neurons via a protein-protein dependent mechanism. FMRP is known to interact with Ca^2+^-activated and Na^+^-activated K^+^ channels and h-channels via protein-protein interactions (Brown et al., 2010; Deng et al., 2013; 2019; Brandalise et al., 2020). Two potential mechanisms are increasing the number of functional channels on the somatic membrane and increasing the single channel conductance. Indeed, there is evidence that FMRP can regulate K^+^ channel function through either of these mechanisms. Application of FMRP_1-298_ to the intracellular domain of Na^+^ activated K^+^ channels decreases the number of sub-conductance states thus increasing the overall flow through the channel (Brown et al., 2010). In CA3 pyramidal neurons, the presence of FMRP increases Ca^2+^ activated BK currents by interacting with the β4 auxiliary subunit of BK channels (Deng et al., 2013).

We previously showed that I_KA_ is higher and I_KD_ is lower in the soma of ET neurons of FX mice (Kalmbach et al., 2015). Neither viral expression of FMRP nor perfusion of FMRP_1-298_ had any effect on I_KA_ in ET neurons suggesting that unlike D-type K^+^ channels, A-type K^+^ channels are not regulated in a cell autonomous protein-protein dependent manner. Furthermore, it suggests that changes in A-type K^+^ channel function in ET neurons of the prefrontal cortex may occur early in development and persist into adulthood. Future experiments investigating the developmental time course of changes in voltage-gated ion channels would be particularly revealing as they could identify a critical window for therapeutic intervention.

## ACKNOWLEDGMENTS

This work supported by National Institutes of Health Grants R01 MH100510 (D.H.B.) The authors wish to thank William Taylor and Eedann McCord for assistance with stereotaxic surgery.

## AUTHOR CONTRIBUTIONS

B.E.K., and D.H.B. designed the study, carried out and analyzed electrophysiology experiments; interpreted results and wrote the manuscript.

## REFERENCES

Brandalise F, Kalmbach BE, Mehta P, Thornton O, Johnston D, Zemelman BV, Brager DH. Fragile X Mental Retardation Protein bidirectionally controls dendritic Ih in a cell-type specific manner between mouse hippocampus and prefrontal cortex. J Neurosci (May 28, 2020). doi: 10.1523/JNEUROSCI.1670-19.2020.

Brown MR, Kronengold J, Gazula V-R, Chen Y, Strumbos JG, Sigworth FJ, Navaratnam D, Kaczmarek LK. Fragile X mental retardation protein controls gating of the sodium-activated potassium channel Slack. Nat Neurosci 13: 819–821, 2010.

Castellino RC, Morales MJ, Strauss HC, Rasmusson RL. Time- and voltage-dependent modulation of a Kv1.4 channel by a beta-subunit (Kv beta 3) cloned from ferret ventricle. Am J Physiol 269: H385–91, 1995.

Darnell JC, Van Driesche SJ, Zhang C, Hung KYS, Mele A, Fraser CE, Stone EF, Chen C, Fak JJ, Chi SW, Licatalosi DD, Richter JD, Darnell RB. FMRP stalls ribosomal translocation on mRNAs linked to synaptic function and autism. Cell 146: 247–261, 2011.

Dembrow NC, Chitwood RA, Johnston D. Projection-specific neuromodulation of medial prefrontal cortex neurons. Journal of Neuroscience 30: 16922–16937, 2010.

Deng P-Y, Carlin D, Oh YM, Myrick LK, Warren ST, Cavalli V, Klyachko VA. Voltage-Independent SK-Channel Dysfunction Causes Neuronal Hyperexcitability in the Hippocampus of Fmr1 Knock-Out Mice. Journal of Neuroscience 39: 28–43, 2019.

Deng P-Y, Rotman Z, Blundon JA, Cho Y, Cui J, Cavalli V, Zakharenko SS, Klyachko VA. FMRP Regulates Neurotransmitter Release and Synaptic Information Transmission by Modulating Action Potential Duration via BK Channels. Neuron 77: 696–711, 2013.

D’Adamo MC, Liantonio A, Conte E, Pessia M, Imbrici P. Ion Channels Involvement in Neurodevelopmental Disorders. Neuroscience (May 2020a). doi: 10.1016/j.neuroscience.2020.05.032.

D’Adamo MC, Liantonio A, Rolland J-F, Pessia M, Imbrici P. Kv1.1 Channelopathies: Pathophysiological Mechanisms and Therapeutic Approaches. IJMS 21, 2020b.

El-Hassar L, Song L, Tan WJT, Large CH, Alvaro G, Santos-Sacchi J, Kaczmarek LK. Modulators of Kv3 Potassium Channels Rescue the Auditory Function of Fragile X Mice. Journal of Neuroscience 39: 4797–4813, 2019.

Gross C, Yao X, Pong DL, Jeromin A, Bassell GJ. Fragile X mental retardation protein regulates protein expression and mRNA translation of the potassium channel Kv4.2. Journal of Neuroscience 31: 5693–5698, 2011.

Gu N, Vervaeke K, Storm JF. BK potassium channels facilitate high-frequency firing and cause early spike frequency adaptation in rat CA1 hippocampal pyramidal cells. J Physiol (Lond) 580: 859–882, 2007.

Guan D, Lee JCF, Higgs MH, Spain WJ, Foehring RC. Functional roles of Kv1 channels in neocortical pyramidal neurons. 97: 1931–1940, 2007.

Guan D, Pathak D, Foehring RC. Functional roles of Kv1-mediated currents in genetically identified subtypes of pyramidal neurons in layer 5 of mouse somatosensory cortex. 120: 394–408, 2018.

Higgs MH, Spain WJ. Kv1 channels control spike threshold dynamics and spike timing in cortical pyramidal neurones. J Physiol (Lond) 589: 5125–5142, 2011.

Jan LY, Jan YN. Voltage-gated potassium channels and the diversity of electrical signalling. J Physiol (Lond) 590: 2591–2599, 2012.

Jerng HH, Pfaffinger PJ, Covarrubias M. Molecular physiology and modulation of somatodendritic A-type potassium channels. Mol Cell Neurosci 27: 343–369, 2004.

Johnston D, Christie BR, Frick A, Gray R, Hoffman DA, Schexnayder LK, Watanabe S, Yuan L-L. Active dendrites, potassium channels and synaptic plasticity. 358: 667–674, 2003.

Kalmbach BE, Johnston D, Brager DH. Cell-Type Specific Channelopathies in the Prefrontal Cortex of the fmr1-/y Mouse Model of Fragile X Syndrome. eNeuro 2, 2015.

Kole MHP, Letzkus JJ, Stuart GJ. Axon initial segment Kv1 channels control axonal action potential waveform and synaptic efficacy. Neuron 55: 633–647, 2007.

Lee HY, Ge W-P, Huang W, He Y, Wang GX, Rowson-Baldwin A, Smith SJ, Jan YN, Jan LY. Bidirectional Regulation of Dendritic Voltage-Gated Potassium Channels by the Fragile X Mental Retardation Protein. Neuron 72: 630–642, 2011.

Noebels JL, Avoli M, Rogawski MA, Olsen RW, Delgado-Escueta AV, Brenner R, Wilcox KS. Potassium Channelopathies of Epilepsy.

Ordemann GJ, Apgar CJ, Brager DH. D-type potassium channels normalize action potential firing between dorsal and ventral CA1 neurons of the mouse hippocampus. Journal of Neurophysiology 121: 983–995, 2019.

Ramos A, Hollingworth D, Adinolfi S, Castets M, Kelly G, Frenkiel TA, Bardoni B, Pastore A. The structure of the N-terminal domain of the fragile X mental retardation protein: a platform for protein-protein interaction. Structure 14: 21–31, 2006.

Routh BN, Johnston D, Brager DH. Loss of Functional A-Type Potassium Channels in the Dendrites of CA1 Pyramidal Neurons from a Mouse Model of Fragile X Syndrome. Journal of Neuroscience 33: 19442–19450, 2013.

Routh BN, Rathour RK, Baumgardner ME, Kalmbach BE, Johnston D, Brager DH. Increased transient Na +conductance and action potential output in layer 2/3 prefrontal cortex neurons of the fmr1 -/ymouse. J Physiol (Lond) 595: 4431–4448, 2017.

Shao LR, Halvorsrud R, Borg-Graham L, Storm JF. The role of BK-type Ca2+-dependent K+ channels in spike broadening during repetitive firing in rat hippocampal pyramidal cells. J Physiol (Lond) 521 Pt 1: 135–146, 1999.

Strumbos JG, Brown MR, Kronengold J, Polley DB, Kaczmarek LK. Fragile X mental retardation protein is required for rapid experience-dependent regulation of the potassium channel Kv3.1b. Journal of Neuroscience 30: 10263–10271, 2010.

Wang LY, Gan L, Forsythe ID, Kaczmarek LK. Contribution of the Kv3.1 potassium channel to high-frequency firing in mouse auditory neurones. J Physiol (Lond) 509 (Pt 1): 183–194, 1998.

Zhang Y, Bonnan A, Bony G, Ferezou I, Pietropaolo S, Ginger M, Sans N, Rossier J, Oostra B, LeMasson G, Frick A. Dendritic channelopathies contribute to neocortical and sensory hyperexcitability in Fmr1(−/y) mice. Nat Neurosci 17: 1701–1709, 2014.

Zhang Y, Brown MR, Hyland C, Chen Y, Kronengold J, Fleming MR, Kohn AB, Moroz LL, Kaczmarek LK. Regulation of neuronal excitability by interaction of fragile X mental retardation protein with slack potassium channels. Journal of Neuroscience 32: 15318–15327, 2012.

